# Cross-generational heritability analysis of physiological traits in *Porites astreoides* across an inshore-offshore gradient in the Lower Florida Keys

**DOI:** 10.1101/2022.03.09.483602

**Authors:** Yingqi Zhang, Shelby J. Barnes, Carly D. Kenkel

**Affiliations:** Department of Biological Sciences, University of Southern California, 3616 Trousdale Parkway, Los Angeles, CA 90089, United States

**Keywords:** coral, life stage, bleaching, thermal tolerance, local adaptation

## Abstract

Estimating the heritable genetic variation in fitness-related traits is key to projecting the adaptive evolution of organisms in response to a changing environment. While heritability studies on reef-building corals to date support adaptive capacity, little is known about the dynamics of trait heritability across life stages in which distinct selective pressures can have long-lasting effects both within and across generations. In this study, we obtained heritability estimates for energetic and thermal stress response traits in larval, recruit, and adult *Porites astreoides* from two populations in the Lower Florida Keys. To induce bleaching phenotypes among individual families, larvae were exposed to a 4-day thermal stress at 32 °C, whereas adults and recruits received the same treatment for 22 days. Origin-dependent tolerance was observed in two life stages where offshore recruits lost more symbiont cells under heat than inshore recruits compared to their respective controls and heat-treated offshore adults suffered a greater loss in total protein content. Surprisingly, larvae appeared to be largely insensitive to heat regardless of origin. Broad sense heritability (*H*^*2*^) estimates varied greatly among traits and life stages, which may reflect changes in the relative importance of genetic and environmental variation throughout development. Notably, more than 80% of the variation in larval chlorophyll a concentration was attributed to genetic factors. The overall moderate to high *H*^*2*^ estimates measured here suggest these corals have considerable potential to adapt to environmental change.

## Introduction

Understanding the adaptive potential of marine organisms is essential to predict population dynamics and persistence in light of rapidly changing ocean temperature and chemistry (Munday et al., 2013; IPCC 2021). For quantitative traits, heritability, which is defined as the proportion of total phenotypic variation attributed to genetic variation, is often used as a metric to estimate the evolvability of those traits under selection (Falconer and Mackay, 1996). Specifically, broad sense heritability (*H*^*2*^) includes all genetic factors, including both additive genetic effects and non-additive genetic effects such as dominance and epistasis. In contrast, narrow sense heritability (*h*^*2*^) is a fraction of *H*^*2*^ and only accounts for additive genetic factors, which are guaranteed to be inherited by the next generation via sexual reproduction. Therefore, *h*^*2*^ describes the true adaptive potential of a given trait but requires knowledge of relatedness or pedigree.

Scleractinian corals are foundation species in reef ecosystems but have undergone severe population declines in recent decades due to increased anthropogenic activities (Hoegh-Guldberg et al., 2007; Moberg & Folke, 1999). The most common stress response in corals is bleaching, which refers to the whitening of tissues resulting from the dissociation of endosymbiotic algae from the coral host (Lesser, 2011). Elevated temperature has been repeatedly shown to be a major cause of mass bleaching events worldwide (Hughes et al., 2018; Jokiel & Coles, 1990). Factors that influence the thermal tolerance of corals include host genetics (Bay & Palumbi, 2014), diversity and flexibility of endosymbiont (family Symbiodiniaceae) (Berkelmans & van Oppen, 2006), and other members of the microbial community (Voolstra et al., 2021). Each component of the coral holobiont is characterized by vastly different generation time (3-74 days for symbiont *in hospite* vs. 4-20 years for coral host) (Muscatine et al., 1984; Babcock, 1991), thus the rate of evolutionary change in each symbiont partner may not align, further complicating the projection of future reefs as sea surface temperature continues to increase.

Similar to other marine invertebrates, corals exhibit complex life history cycles that oscillate between the planktonic larval stage and the benthic juvenile/adult stages (Richmond & Hunter, 1990). Different life stages encounter distinct selective pressures but phenotypic changes in a given life stage resulting from interactions with its environment can have long-lasting effects both within and across generations (Marshall & Morgan, 2011; Putnam, 2021). The majority of work to date has focused on the effects of abiotic stressors on the physiology and critical developmental processes of one distinct life stage and only a handful of studies have compared phenotypic response to similar stress among multiple life stages (Kenke et al., 2015; Putnam et al., 2010; Putnam & Gates, 2015). Even less is known about how variable the heritability of a given trait may be across a coral’s life span (Bairos-Novak et al., 2021). Additive genetic variance (and hence *H*^*2*^) of fitness-related traits has been shown to increase with age in birds (mute swan and collared flycatcher) and terrestrial mammals (red deer and soay sheep), likely explained by the accumulation of deleterious mutations, or positive selection on genes that increase fitness at earlier life stages but have the opposite effects later in life (Wilson et al., 2008). Carlon et al. (2011) documented lower *H*^*2*^ estimates of coral skeletal traits in juvenile *Favia fragum* than in adult colonies and attributed the differences to environmental effects. More work is needed to shed light on the dynamics of heritability over distinct coral life stages to obtain a holistic understanding of the adaptive potential of critical phenotypes, as only focusing on a single life stage can be misleading (Albecker et al. 2021).

In this study, we aimed to quantify thermal performance of *Porites astreoides* at adult, larval, and recruit life stages as well as estimate broad sense heritability of various physiological traits shared across all stages. We utilized two distinct populations originating from an inshore and an offshore reef environment in the Lower Florida Keys. Compared to offshore reefs at similar latitudes, inshore reefs experience greater thermal variation and more extreme summer temperatures (Kenkel et al., 2015; Lirman & Fong, 2007). Nonetheless, inshore coral populations are characterized by greater thermal tolerance, higher cover, higher growth and calcification rate (Kenkel et al., 2013; Manzello et al., 2015; Manzello et al., 2019). Prior studies on inshore and offshore *P. astreoides* populations revealed distinct host genetic structures and transcriptomic profiles but shared symbiont type (ITS2 type A4/A4a), indicating that the animal host might play a bigger role in conferring thermal tolerance (Kenkel et al., 2013; Kenkel & Matz, 2017). We aimed to take advantage of standing genetic variation and origin-specific thermal traits between the two populations to answer the following two questions: 1. Are inshore larvae and recruits also more heat resistant than the offshore offspring? 2. Are traits associated with thermal tolerance more or less heritable depending on life stage?

## Methods

### Spawning collection and larval thermal stress experiment

A total of 30 adult *Porites astreoides* colonies were collected from an inshore reef site (Summerland Shoals Patch: 24° 36.346′ N, 81° 25.742′ W, n=15) and an offshore reef site (Big Pine Ledges: 24° 33.174′ N, 81° 22.809′ W, n=15) five days before the new moon on April 29, 2019 under permit #FKNMS-2018-033. Colonies were transported to Mote Marine Laboratory’s Elizabeth Moore International Center for Coral Reef Research and Restoration and kept in a shaded (70% PAR reducing) raceway to monitor for planulation. Plastic numerical tags were attached to the bottom of colonies using marine epoxy putty (All-Fix) to track individual identities. Colonies were then placed into flow-through larval collection chambers before sunset according to Kuffner et al. (2006). Beaker traps were checked for larvae the following morning (April 30, 2019). Most inshore colonies produced enough larvae to be included in the subsequent experiment while only one offshore colony (Family 34) had sufficient planulation. As a result, collection chambers were deployed for a second night to capture more spawning colonies that originated from the offshore site.

A subset of larvae from 5 inshore and 5 offshore colonies was sampled (n=3×10 per family) and frozen at −80 °C prior to analyzing a suite of physiological assays. Another subset of larvae with the same replication were then subjected to a moderate heat stress challenge beginning on May 1 following the same methods described in Zhang et al. (2019) (Fig. 1). Briefly, three replicate plastic bins were set up for both control (26 °C) and treatment (32 °C) groups. Treatment was set to 32 °C because inshore reefs commonly experience this temperature during late summer months (Aug-Sept) when bleaching is likely to occur (Kenkel et al., 2015). Additionally, 32 °C induces a level of stress that can produce distinct bleaching responses among individuals (Zhang et al., 2019). Each bin was filled with 30 L seawater and equipped with a SL381 submersible water pump, a 100 W aquarium heater, and a HOBO temperature logger (Onset). One group of 10 larvae per spawning colony was added to a 70 uM cell strainer (Grenier Bio-One) which served as a floating netwell in each replicate bin (n=3×10 larvae per family per temperature). Target temperature for the heat treatment bins was achieved by increasing the temperature by 0.5 °C per hour over a 12-hour window and maintained for 4 days. At the end of the exposure period, swimming larvae in each netwell were sampled (with seawater removed) and frozen at −20 °C for subsequent physiological assays.

**Fig. 1.**
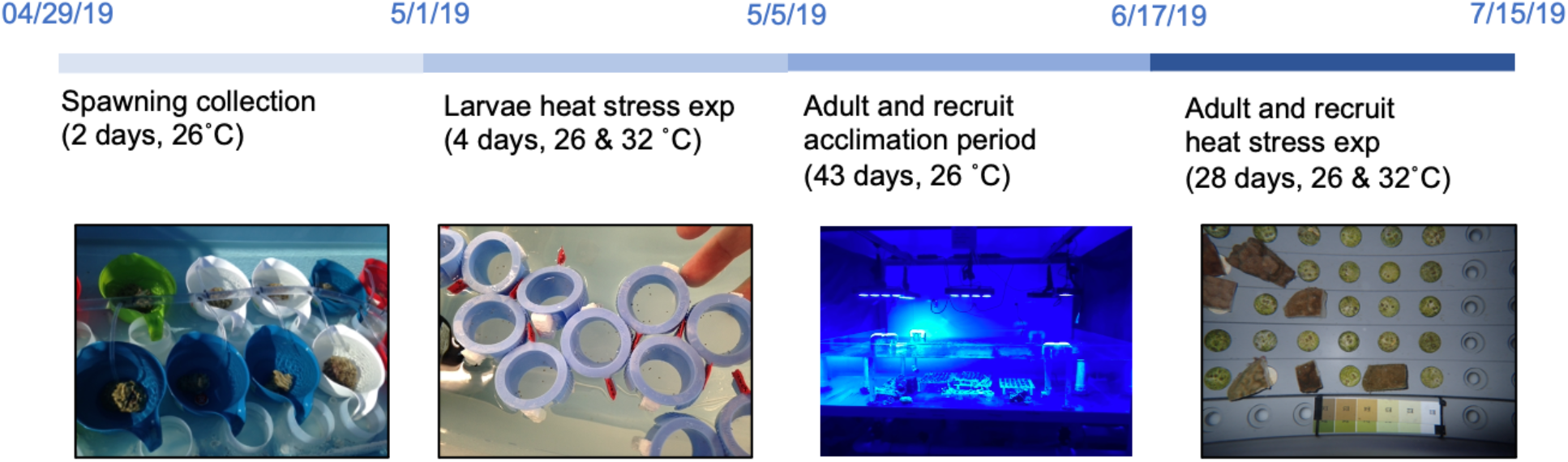
Experimental design and timeline including ten *Porites astreoides* families

Remaining larvae were settled onto aragonite plugs preconditioned with crustose coralline algae (CCA) by family, with the goal of recruiting between 3 and 4 individuals on the top side of each plug to avoid fusion of recruits. For each larval family, 19 plugs were secured by two staggered egg crates placed in a 1.2 L plastic food storage container. After adding seawater to the containers, larvae were gently pipetted onto the plugs and into the water column. Containers were monitored daily and water changes were performed every two days, after which new larvae were introduced to facilitate further recruitment. To securely transport recruits to the University of Southern California (USC), plugs with ideal settlement densities were superglued to the caps of 50 mL falcon tubes, which were then filled up with seawater to displace any air bubbles. Adult colonies that produced larvae were wrapped in wet bubble wrap and transported to USC along with the recruits.

### Adult and recruit thermal stress experiments

Adult coral fragments and their recruits were allowed a 37-day recovery and acclimation period in a 500 L holding tank at the Cnidarian Evolutionary Ecology Lab aquarium room at USC (26 °C, 80 μmol photons m^−2^ s^−1^, 5 cm s^−1^). After a month of recovery (June 5), six ca. 15 cm^2^ fragments were obtained from each adult colony and mounted on clean aragonite plugs using superglue. Adult fragments were then returned to the acclimation tank. To quantify the thermal tolerance of adult and recruit life stages, three replicate 50 L tanks were set up for both control (26 °C) and treatment (32 °C) groups. On June 13, 2019, one adult fragment per colony and an average of 8.5 recruits per family were randomly assigned to each tank. Recruit numbers as well as settlement patterns varied across families, thus each tank received between 1 and 6 plug(s) to achieve an even distribution of recruits across tanks. The experimental tanks were individually outfitted with submersible water pumps (Sicce Syncra Silent 0.5, 185 GPH), 150W HMA-S thermal-regulated heaters (Finnex), and AI Prime HD lights (Aqua Illumination) programmed to imitate the light cycle in FL, with an average daytime PAR of 85-115 μmol photons m^−2^ s^−1^ reaching the tank bottom. Adult and recruit plugs were acclimated to their individual tank conditions with recirculating flow for 4 days. Adult fragments and recruit plugs were cleaned using a small brush every two days for the duration of the experiment to remove any filamentous algal growth. The thermal stress challenge started on June 17, when recirculating flow was turned off for each tank to prevent pseudo replication. Temperature in the individual heat treatment tanks was slowly ramped to 32 °C over 6 days (1 °C increase per day) and maintained for 22 more days until July 15. HOBO temperature loggers were deployed to track the temperature profile throughout the exposure. A 10% water change was performed every week and the concentrations of potential nitrogen metabolic wastes (including NH_4_^+^, NO_3_^−^, and NO_2_^−^) were measured daily using commercial aquarium test kits. Salinity was maintained at 35 ppt by topping off with distilled water (300-500 ml for control tanks and 500-1300 ml for heat tanks per day).

To track bleaching status and recruit growth over the course of the thermal stress exposure, photographs were taken under identical illumination using an Olympus Tough camera TG-4/TG-5 (Olympus America Inc.) at the initial timepoint (T_0_), final timepoint (T_28_), and 6 additional time points in between (T_7_, T_11_, T_14_, T_18_, T_21_, T_25_). Starting with the control group, PVC racks were removed from each tank and immediately placed into a Nally bin filled with seawater. To minimize the warping effect along the edges of the photos, racks were divided into 4 quadrants and each quadrant was photographed individually to make sure the corals were centered (Figure. 1). A subset of the Coral Health Chart (Siebeck et al., 2006), E1-E6 and B1-B6, was ziptiped to an empty rack that was placed against the coral racks to serve as a color and size reference. Each quadrant was photographed three times with the camera lens just submerged. After all four quadrants were covered, the coral racks were immediately transferred back to their respective treatment tank. The Nally bin was then heated to 32 °C before corals from the heat treatment group were transferred and photographed.

After 22 days of exposure, recruits were scraped off the plugs using a razor blade and pooled into 1.5 ml Eppendorf tubes by family per replicate tank. Adult fragments were removed from their plugs and wrapped in pre-labeled tin foil. All samples were frozen immediately at −80 °C for subsequent physiological assays.

### Physiological assays

Similar to Zhang et al. (2019), larvae (3 replicates of 10 larvae per family) and recruit (3 replicates of multiple recruits per family, see Table S1) samples were thawed on ice and mixed with 100 μl extraction buffer (50 mM phosphate buffer, pH 7.8, with 0.05 mM dithiothreitol). The mixture was further homogenized by back pipetting to free symbiont cells from host tissues and the final volume of the homogenate was recorded to account for residual seawater. For recruit samples, skeletal debris was allowed to settle for a minute and the aqueous tissue layer was further transferred to a new set of tubes. To quantify symbiont cell density, 20 μl of the homogenate was fixed in 20 μl of a 20% formalin solution (10% final concentration) and triplicate 10 μl aliquots were assessed under a compound microscope at 100x magnification using a hemocytometer. The remaining slurry was centrifuged for 3 min at 1500 x g at 4 °C to pellet symbiont cells for chlorophyll quantification and isolate host tissue supernatant for protein analysis. Symbiont cell pellets were resuspended in 90% acetone and further broken down by shaking with metal beads in a TissueLyser II (Qiagen) for 90 s. After an overnight incubation at −20 °C, the solution was centrifuged for 5 min at 10,000 x g at 4 °C. Triplicate 50 μl aliquots of the resultant supernatant were measured for absorbance at 630, 647, and 664 nm using Synergy H1 microplate reader (Biotek). Chlorophyll a concentration was determined using the equation specified in Ritchie (2008). Soluble host protein was measured colorimetrically using triplicate 10 ul aliquots of the host supernatant with the RED 660™ protein assay kit (G-Biosciences) following the manufacturer’s instructions. Measured concentrations were multiplied by initial homogenate volume to account for differential dilution. Larval traits were normalized by number of larvae, whereas recruit traits were normalized by total recruit surface area (see details below) as individual recruit size was quite variable.

The same workflow also applied to adult sample processing with a few modifications: each thawed adult fragment was mixed with 10 ml extraction buffer in a plastic zipper bag (Plymor) over ice and airbrushed to remove the tissue, which was then homogenized by a tissue homogenizer (VWR^®^ Model 200) for 1 min following (Palmer et al., 2010) and the final volume was recorded. Symbiont cells were fixed by mixing 250 μl of homogenate with 125 μl 20% formalin solution (6.7% final concentration). One ml of homogenate was aliquoted to conduct chlorophyll analysis. Host tissue was isolated by centrifuging the remaining homogenate (ca. 8.75 ml) at 1500 xg. Skeletons were cleaned using 10% bleach and air dried. Surface area was assessed using the single wax dipping method (Veal et al., 2010) and was used to standardize adult traits.

Adult and recruit photos were analyzed in ImageJ (Schneider et al., 2012). To assess visual bleaching, color scores were assigned to individuals at each time point following Siebeck et al. (2006). Briefly, the Coral Health Chart color reference was used to generate a standard curve of mean grayness values (or an average of R, G, B values) from the standardized color scores (D1-D6) within each photo. The entire surface area of recruits was traced and three unshaded subsets of adult fragments were selected to measure mean grayness value, which was then converted to a color score using the standard curve. A recruit was considered dead (recorded as NA) if no visible tissue remained. Recruit size was determined by tracing the outline of each individual and recording the pixel area. A line of known distance on the Coral Health Chart was used as a reference to convert pixel area to metric area. Growth of individual recruits was calculated as the difference in size between initial and final timepoints.

### Statistical analyses

All statistical analyses were conducted in R 4.0.3 (R Core Team, 2020). Linear mixed effects models (*lme4* package, Bates et al., 2015) were used to analyze all physiological traits, including symbiont density, chlorophyll a concentration, total protein content, change in color score, and recruit growth. Treatment (levels: control and heat) and origin (levels: inshore and offshore) were included as fixed effects. Family and tank (for adults and recruits only) were included as random effects. Models were evaluated for normality and homoscedasticity using diagnostic plots. Log-transformation was performed on trait data that did not satisfy the model assumptions of normality and absence of heteroscedasticity of residuals. The null hypothesis was rejected at an alpha of 0.05. Correlation coefficients were calculated for traits that were shared between life stages to investigate the strength of familial effect across stages and values were displayed in matrices using the *corrplot* package (Wei et al., 2017). Trait data were grouped by family and treatment and thus each comparison consisted of 20 data points.

Broad sense heritability (*H*^*2*^) of measured traits was estimated using MCMC models (*MCMCglmm* package, Hadfield, 2010) for each life stage, including treatment and origin as fixed effects and family as a random effect. Traits were log-transformed as necessary to meet the normality requirement. All models were run for 100,000 iterations, with the first 10,000 discarded to ensure convergence and every 20 subsequent parameter values sampled to minimize autocorrelation. The resulting effective sample size was 4,500. *H*^*2*^ values were calculated as the ratio of variance attributable to the random familial effect over total variance.

## Results

Adults, larvae, and recruits all experienced a significant decline in symbiont density in response to heat stress (*p* < 0.05, Figure 2a–c). The response also differed by population in the early life stages, where recruits originating from inshore families lost fewer symbionts than offshore recruits, by 5% on average (*t* = −2.86, *df* = 48, *p* < 0.01, Figures 2c) while inshore larvae tended to lose more symbionts than offshore larvae post treatment (*p* = 0.056, Figures 2b). Inshore larvae in general contained 50% more symbiont cells than offshore larvae (*t* = −3.35, *df* = 8, *p* = 0.01, Figures 2b). Mean adult symbiont density correlated positively with mean recruit symbiont density across families (*r* = 0.66, *p* < 0.01) but no correlations with larvae were apparent (Figure S1a).

**Fig. 2.**
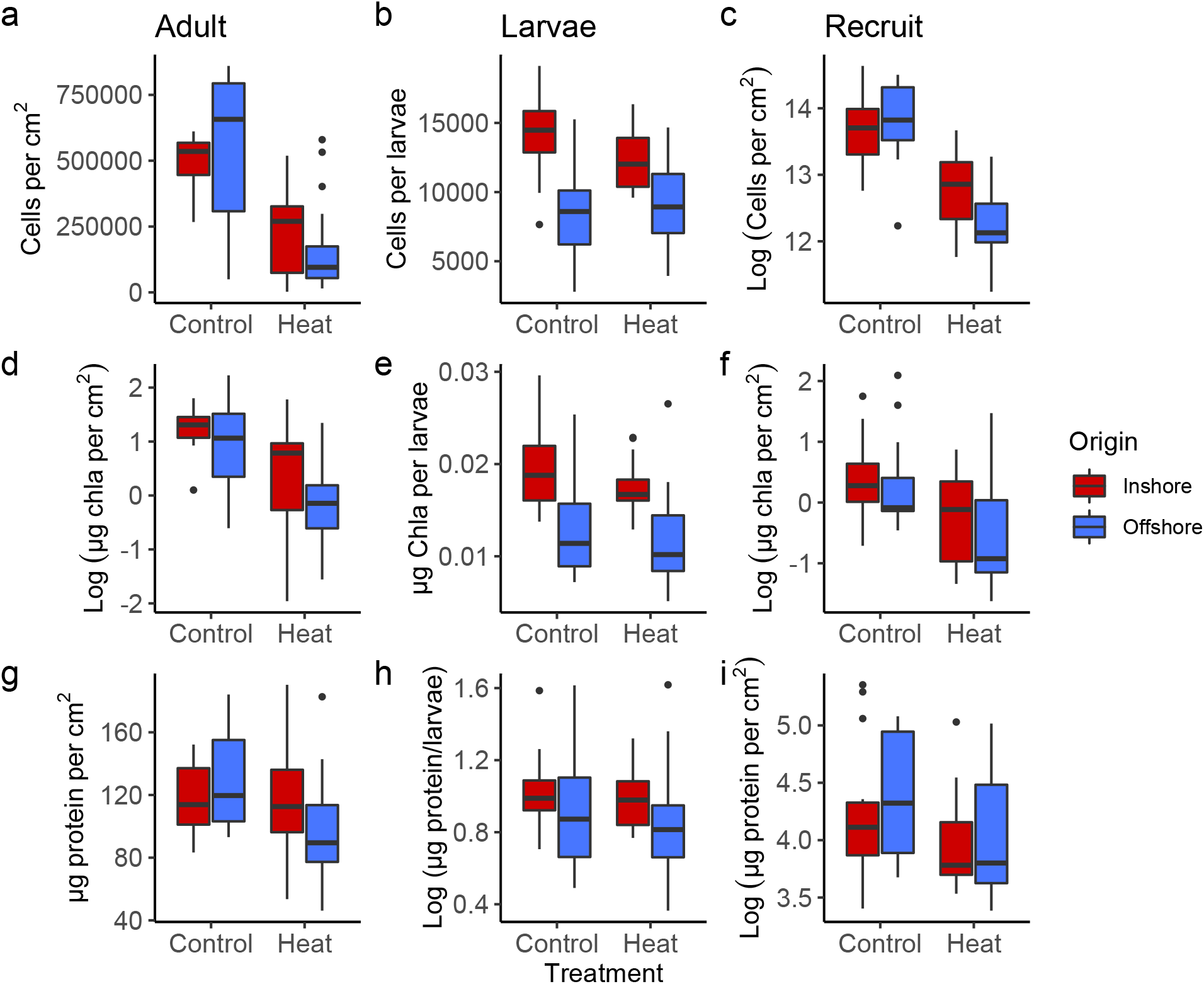
Standardized physiological parameters (mean ± SEM) to experimental conditions shared across all life stages separated by reef origin. Each column represents a distinct life stage (left to right: adult, larvae, and recruit) and each row represents a distinct trait (top to bottom: symbiont density, chlorophyll a concentration, total soluble protein content). Values were log-transformed if not normally distributed

Chlorophyll a concentration declined by 85% (*t* = −3.86, *df* = 48, *p* < 0.001, Figure 2d) in heat-treated adults and tended to decrease (*t* = −1.97, *df* = 45, *p* = 0.055, Figure 2f) in heat-treated recruits compared to their respective controls. Larval chlorophyll a differed significantly by origin where inshore larvae had 49% (*t* = −2.60, *df* = 8, *p* < 0.05, Figure 2e) more chlorophyll a than offshore larvae. Positive correlations were detected between chlorophyll a concentration in adults and larvae as well as between adults and recruits (*r* = 0.46 and 0.45 respectively, *p* < 0.05), but concentrations in recruits and larvae were unrelated (Figure S1b).

No fixed effect of origin or treatment was observed for total soluble host protein content for any life stage, save for a marginal treatment effect in recruits where protein content tended to be reduced in response to heat stress (*p* = 0.059, Figure 2i). However, a significant interaction was detected in adults where offshore coral lost 23% more protein after the heat exposure period compared to inshore coral (*t* = - 2.18, *df* = 48, *p* < 0.05, Figure 2g). No significant correlations were found between any two life stages in terms of protein content (Figure S1c).

The standardized color score of adult and recruit corals was on average lower at the end of the experiment than when it started, with heat-treated individuals experiencing a greater degree of paling, by more than 2-fold in both life stages (*p* < 0.05, Figure S2). The correlation between change in color score over time in adults and recruits was not significant. Individual recruit size also decreased after the exposure period, but recruits from the heat group suffered less of a decrease than those from the control group (*t* = 3.05, *df* = 48, *p* < 0.01, Figure S3).

Significant heritability estimates were detected for multiple physiological traits but varied greatly across life stages. The average broad sense heritability estimate for symbiont density was highest in the adult stage (*H*^*2*^= 0.65, 95% confidence interval (CI): 0.27, 0.96), followed by larval (*H*^*2*^= 0.54, 95% CI: 0.25, 0.84) and recruit stages (*H*^*2*^= 0.32, 95% CI: 0.0009, 0.61) (Figure 3). The highest *H*^*2*^ value was detected for chlorophyll a concentration in larvae (*H*^*2*^= 0.84, 95% CI: 0.70, 0.96), indicating more than 80% of the variance in chlorophyll a was explained by genetic factors, which include additive genetic effects, epistasis, and maternal effects. In comparison, the *H*^*2*^ estimate for chlorophyll a yielded a lower value of 0.53 (95% CI: 0.27, 0.84) in adults and a much lower value of 0.07 (95% CI: 3×10^−5^, 0.22) in recruits. Heritability estimates for protein content exhibited the opposite trend where recruits had the highest value of 0.67 (95% CI: 0.42, 0.92), followed by larvae (*H*^*2*^= 0.61, 95% CI: 0.33, 0.87) and adults (*H*^*2*^= 0.26, 95% CI: 3×10^−7^, 0.58). The *H*^*2*^ of change in color score between final and initial time points was 0.35 (95% CI: 0.03, 0.66) for adults and 0.11 (95% CI: 0.01, 0.25) for recruits.

**Fig. 3.**
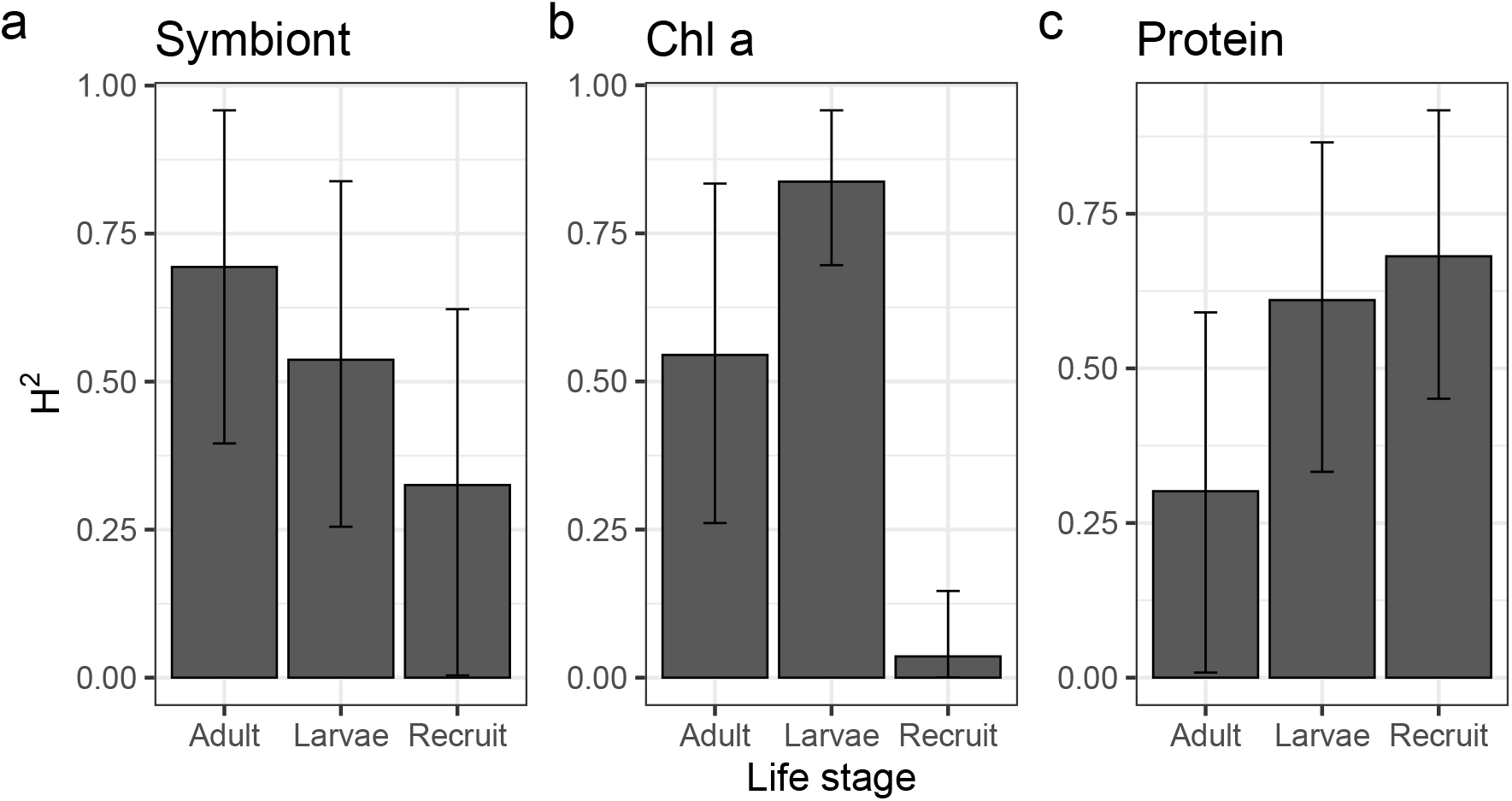
Broad-sense heritability estimate (mean ± 95% C.I.) for three shared physiological traits at different life stages

## Discussion

Heat induced a significant bleaching response in all three life stages, as indicated by the reduction in symbiont density and/or chlorophyll a concentration after treatment (Figure 2a–f). Adult corals and their recruits in the 32 °C group also lost more pigmentation as indicated by larger decreases in color score (Figure S2). Interestingly, larvae appeared to be more heat resistant compared to adults and recruits. In particular, symbiont density was unaffected by treatment in several larval families (5, 11, 38, 39) and even slightly elevated in two offshore families 32 and 34 (Figure S4b). No treatment effect was detected at all for larval chlorophyll concentration (Figure 2e). One possible explanation for the muted response to heat in larvae is a shorter exposure period (i.e., 4 days as opposed to 22 days for adults and recruits). From a developmental standpoint, it would be difficult to extend the exposure window as *P. astreoides* larvae become competent quickly after release (Ritson-Williams et al., 2016) and high temperature tends to expedite settlement (Nozawa & Harrison, 2005). Another possibility may be that larvae are indeed more tolerant of thermal stress than later life stages, although prior evidence has been mixed and studies emphasizing both the vulnerability and robustness of coral larvae can be found. Earlier studies showed that larval survivorship and settlement in broadcast spawning coral species *Diploria strigosa* and *Acropora palmata* were significantly reduced when subjected to sublethal thermal stress (30-32 °C) (Bassim & Sammarco, 2003; Randall & Szmant, 2009). Symbiotic larvae of *Pocillopora damicornis*, a brooding coral like *P. astreoides,* suffered from impaired metabolism under a combination of high temperature and CO_2_ stresses (Rivest & Hofmann, 2014). Another study on the same species reported an over 50% reduction in dark-adapted maximum quantum yield of photosystem II of heat-exposed larvae compared to adults under the same treatment (Putnam et al., 2010). Conversely, our previous study highlighted incredible thermal tolerance of *P. astreoides* larvae where significant mortality under 36 °C was only observed after >24 hours of exposure (Zhang et al., 2019), mirroring the survival data for aposymbiotic *Acropora millepora* larvae subjected to a similar level of stress (Dixon et al., 2015). *A. millepora* larvae also survived well and underwent normal development after a 5-day exposure to 32 °C (Meyer et al., 2009). Larval thermal tolerance is therefore likely species- and phenotype-specific. More cross-generational or even multi-generational studies are needed to clarify these conflicting results. Ultimately, the performance of larvae also needs to be contextualized in a wider developmental framework, as short-term sublethal thermal stress experienced by larvae can negatively affect recruitment and post-recruitment survival (Edmunds et al., 2001; Ross et al., 2013).

Population-specific responses to heat stress were observed in both adults and recruits. Notably, inshore adult corals lost less protein content and inshore recruits lost fewer symbiont cells respectively in comparison to their offshore counterparts (Figure 2g and 2c). Adult *P. astreoides* colonies from the inshore environment in the Lower Florida Keys have been repeatedly shown to exhibit higher bleaching tolerance based on multiple phenotypes, including bleaching color score, brightness in the red channel, and photochemical yield of the algal endosymbiont (Kenkel et al., 2013; Kenkel et al., 2015). In this study, surprisingly, inshore adults failed to outperform the offshore adults in response to heat based on the two measured bleaching phenotypes, although similar trends were apparent. This could be explained by the comparatively smaller sample size and noticeable variation among individual colonies; families 11 and 17 were more similar to offshore families whereas family 39 was more inshore-like (Figure S4a). Although among-family variation in thermal tolerance of *P. astreoides* adults originating from the same location has not been investigated before, our prior study documented significant family effects on larval survival and physiology, where a greater percentage of phenotypic variation was due to parental colony identity than other factors such as day of release and reef origin (Zhang et al., 2019). In addition, the significant positive correlation in symbiont density between individual adult and recruit families suggest strong family effects can persist into the following life stage (Figure S1a). Most importantly, the maintenance of symbiont densities under elevated temperature presents the first evidence of enhanced bleaching tolerance in inshore offspring. This may indicate that the differences in tolerance between inshore and offshore recruits has a heritable basis given that these recruits lacked prior exposure to different thermal regimes, which is further supported by the analysis of heritability.

Reduction in recruit size was likely due to lack of feeding throughout the experimental period (Figure S3). Feeding *Pocillopora acuta* with brine shrimp (*Artemia* spp.) three times a week doubled colony growth and increased symbiont density as well as their maximum quantum yield (Huang et al., 2020). We chose not to feed the corals mainly due to the concern of increased fouling in a small, closed system. More importantly, neither treatment group was provisioned with external food sources, so it is still valid to make conclusions on population level differences in heat tolerance by comparing control and heat-treated corals, although results may differ under replete conditions. Corals derive the majority of their daily energetic requirements from symbiotic Symbiodiniaceae (Muscatine and Porter, 1977). However, certain species rely more heavily on heterotrophic feeding during bleaching events when their autotrophic source is limited (Grottoli et al., 2006). It is unclear whether our focal species *P. astreoides* has flexible heterotrophic capacities and therefore hard to predict the actual effects of feeding. Based on the limited trophic plasticity in its Pacific congeners *P. compressa* and *P. lobata* (Grottoli et al., 2006), feeding might have a minor role in mitigating the effect of bleaching in *P. astreoides.*

In general, we found moderate (*H*^*2*^= 0.25~0.50) to high (*H*^*2*^> 0.50) broad sense heritability estimates across all traits examined (Figure 3). This finding reinforces the main conclusion of a recent meta-analysis by Bairos-Novak et al. (2021) which found that the majority of coral physiological traits (with few exceptions such as gene expression and photochemistry) exhibited relatively high heritability. For instance, our *H*^*2*^ estimate of adult symbiont density (0.65) is on par with an estimate of 0.71 for the same trait in adult *Orbicella faveolata* during a natural bleaching event (Manzello et al., 2019). These findings indicate adaptive capacity, collectively projecting a positive outlook for the survival of coral communities under changing environmental conditions. One important caveat, however, is that the heritability values derived from this study likely overestimate the true adaptive potential of our focal traits because *h*^*2*^ is only a subset of *H*^*2*^ (Falconer and Mackay, 1996). Nonetheless, *h*^*2*^ for highly heritable traits in certain life stages, for instance, chlorophyll a in larvae and protein in recruits, is expected to be significant as well, as *h*^*2*^ is typically only 1.4-2.5 fold lower than *H*^*2*^ (Bairos-Novak et al., 2021). Another factor that may bias estimates is the assumption that each spawning colony possesses a unique genetic identity. Given the rapid propensity for settlement in *P. astreoides* larvae, they are expected to have short dispersal potential and therefore high local retention (Jones et al., 2009). It is likely that clones or sibling clusters group over a small spatial scale as larvae of other brooding species have been documented to settle within meters from their parents (Carlon & Olson, 1993).If phenotypically similar colonies were clones and their larval/recruit families were siblings, true heritability values could possibly be higher than our estimates. Conversely, true heritability for a given trait may be lower than estimated if phenotypically dissimilar individuals shared the same genetic identity. In the absence of genotyping data we are unable to rule out the possibility of clones, however, it is unlikely that we collected clonal or sibling adult colonies because we sampled individuals that were at least 10 m apart in the field, a greater distance compared to 1 m used in other studies (Riquet et al., 2021; Serrano et al., 2016).

Despite overall moderate to high heritability levels, estimates differed among life stages and across different physiological traits within a given stage. Although the confidence interval associated with most heritability values is not trivial, it is comparable to similar error estimates in previous analyses (Jury et al., 2019; Kenkel et al., 2015) and in future work can be potentially reduced by larger sample sizes and increased replication. Symbiont density tended to be more heritable in the later adult stage, followed by larvae and recruits (Figure 3). Similarly, recruits had the lowest heritability estimate for chlorophyll a concentration (*H*^*2*^=0.07), whereas over half of the variation in adults and larvae was attributed to genetics. In various vertebrate species, older age is usually associated with high genetic variance in fitness-related traits due to the accumulation of somatic mutation over time (Wilson et al., 2008). A recent study estimated that the table coral *Acropora hyacinthus* accumulates mutations at a similar rate (2.6 mutations per gigabase per year in its coding region) as human somatic cell lines (López & Palumbi, 2020). Given this parallel, it is reasonable to expect increased mutations and genetic variation in older corals with bigger colony sizes. Moreover, those potentially deleterious mutations are suspected to be purged before gamete production (Orive, 2001), which likely contributes to the relatively lower genetic variation in the offspring. However, the rank order of *H*^*2*^ estimates is completely reversed for soluble protein content, where higher values were observed in larvae and recruits rather than adults (Figure 3). Based on an interspecific comparison, the genetic component of protein content was about four times higher in *P. astreoides* larvae (*H*^*2*^= 0.60) than in *A. millepora* adults (*H*^*2*^= 0.15) (Bairos-Novak et al., 2021). One possible explanation is that maternal effects may be dominant during early life stages as energetic reserves are maternally provisioned (Richmond & Hunter, 1990), thus inflating the overall estimate of genetic effects. Moreover, brooding species like *P. astreoides* likely experience larger maternal effects since brooded larvae are subject to maternal environment during early development and inherit symbionts directly from the mother (Richmond & Hunter, 1990). Interestingly, despite the general expectation that maternal effects attenuate over time (Dufty et al., 2002), the estimate of heritability of protein content was similar between recruits and larvae. Studies on small mammals and birds have found that phenotypic variance components constantly fluctuate across developmental stages, resulting in divergent heritability estimates depending on sampling time (Atchley, 1984.; Bourret et al., 2017). Additive genetic variance (*V*_*A*_), in particular, can be modified by relative expression at specific loci and/or the phenotypic effects of those loci especially during early ontology (Atchley, 1984). Indeed, loci involved in metabolism and proteolysis were differentially expressed in *A. millepora* planula larvae and newly-settled polyps (Hayward et al., 2011). It is also possible that different life stages in *P. astreoides* use different sets of genes to respond to environmental stimuli (Ruggeri et al. *in prep*), which may further contribute to the difference in *V*_*A*_ observed between larvae and recruits.

In summary, we observed population-level differences in the response to heat stress in both adult and offspring generations of *P. astreoides* derived from two distinct reef zones in the Lower Florida Keys, highlighting the potential contribution of genetic adaptation and physiological acclimatization to thermal tolerance. Larvae tended to be more bleaching tolerant than adults and recruits from the same lineages, although more studies involving multiple life stages need to be conducted to validate this hypothesis. It is also important to identify an appropriate level and duration of heat stress for each life stage so that responses are comparable. Moreover, we also find that the broad sense heritability of a given trait is widely divergent among different life stages. Two bleaching phenotypes, symbiont density and chlorophyll a concentration, were moderately to highly heritable in adults and larvae, whereas protein content was moderately to highly heritable in larvae and recruits, which likely reflects the fluctuating dynamics between genetic variation and environmental variation as organisms undergo different developmental phases. The overall significant heritability levels of bleaching and nutrient associated traits bode well for the local *P. astreoides* populations as they may be capable of keeping up with the rapidly changing ocean temperatures through adaptation across generations.

## Acknowledgment

We would like to thank E. Bartels and other staff members at Mote Marine Laboratory’s International Center for Coral Reef Research and Restoration for their amazing field and logistical support, W. Million and E. Aguirre for their assistance with field sampling and conducting the experiments, C. Timmons for her help with aquarium maintenance, S. O’Donnell for her dedication to color score analysis. Lastly, we greatly appreciate the Manahan lab (especially M. DellaTorre) at USC for lipid protocol training and troubleshooting despite unpublishable results. This research was supported by start-up funds from the University of Southern California to C. Kenkel.

## Author’s Contributions

YZ and CDK designed the study and performed field collection and larval experiments. YZ and SJB conducted adult and recruit experiments and end sampling. SJB processed adult samples and YZ finished the remaining samples and analysis. YZ wrote the first draft of the manuscript. All authors reviewed, approved of and are accountable for this submission.

## Data Availability

Raw data and R scripts for statistical analysis are available at https://github.com/yingqizhang/Porites2022.

## Conflict of interest

The authors declare no conflict of interest.

## Notes

### Competing Interest Statement

The authors have declared no competing interest.

