## Supplemental Table S1 and Figures S1-S4 for "Cross-generational heritability analysis of physiological traits in *Porites astreoides* across an inshore-offshore gradient in the Lower Florida Keys"

### Supplementary Materials

Table S1
Figures S1-S4

#### Cross-generational heritability analysis of physiological traits in *Porites astreoides* across an inshore-offshore gradient in the Lower Florida Keys

Yingqi Zhang[1*], Shelby J. Barnes[1], Carly D. Kenkel[1]

[1] Department of Biological Sciences, University of Southern California, 3616 Trousdale Parkway, Los Angeles, CA 90089, United States

**Table S1.** Number of recruits in each technical replication.

| Trmt | Rep tank | Family | Num ($T_{initial}$) | Num ($T_{final}$) | Trmt | Rep tank | Family | Num ($T_{initial}$) | Num ($T_{final}$) |
|---|---|---|---|---|---|---|---|---|---|
| Control | 1 | 5 | 15 | 15 | Heat | 1 | 5 | 9 | 9 |
| | | 11 | 8 | 4 | | | 11 | 4 | 1 |
| | | 17 | 8 | 8 | | | 17 | 8 | 8 |
| | | 19 | 11 | 11 | | | 19 | 10 | 10 |
| | | 20 | 9 | 9 | | | 20 | 11 | 11 |
| | | 32 | 7 | 7 | | | 32 | 10 | 10 |
| | | 34 | 8 | 8 | | | 34 | 12 | 12 |
| | | 38 | 7 | 3 | | | 38 | 8 | 7 |
| | | 39 | 7 | 5 | | | 39 | 7 | 7 |
| | | 46 | 10 | 10 | | | 46 | 10 | 9 |
| Control | 2 | 5 | 12 | 12 | Heat | 2 | 5 | 6 | 6 |
| | | 11 | 3 | 2 | | | 11 | 5 | 3 |
| | | 17 | 8 | 8 | | | 17 | 8 | 8 |
| | | 19 | 7 | 7 | | | 19 | 9 | 9 |
| | | 20 | 16 | 16 | | | 20 | 7 | 7 |
| | | 32 | 10 | 10 | | | 32 | 14 | 14 |
| | | 34 | 11 | 11 | | | 34 | 8 | 7 |
| | | 38 | 8 | 7 | | | 38 | 7 | 6 |
| | | 39 | 7 | 5 | | | 39 | 8 | 6 |
| | | 46 | 11 | 11 | | | 46 | 11 | 11 |
| Control | 3 | 5 | 14 | 14 | Heat | 3 | 5 | 11 | 11 |
| | | 11 | 5 | 4 | | | 11 | 5 | 5 |
| | | 17 | 10 | 10 | | | 17 | 8 | 8 |
| | | 19 | 12 | 12 | | | 19 | 12 | 12 |
| | | 20 | 11 | 11 | | | 20 | 10 | 8 |
| | | 32 | 10 | 9 | | | 32 | 15 | 15 |
| | | 34 | 5 | 5 | | | 34 | 12 | 12 |
| | | 38 | 6 | 6 | | | 38 | 10 | 8 |
| | | 39 | 5 | 4 | | | 39 | 6 | 6 |
| | | 46 | 12 | 12 | | | 46 | 12 | 12 |

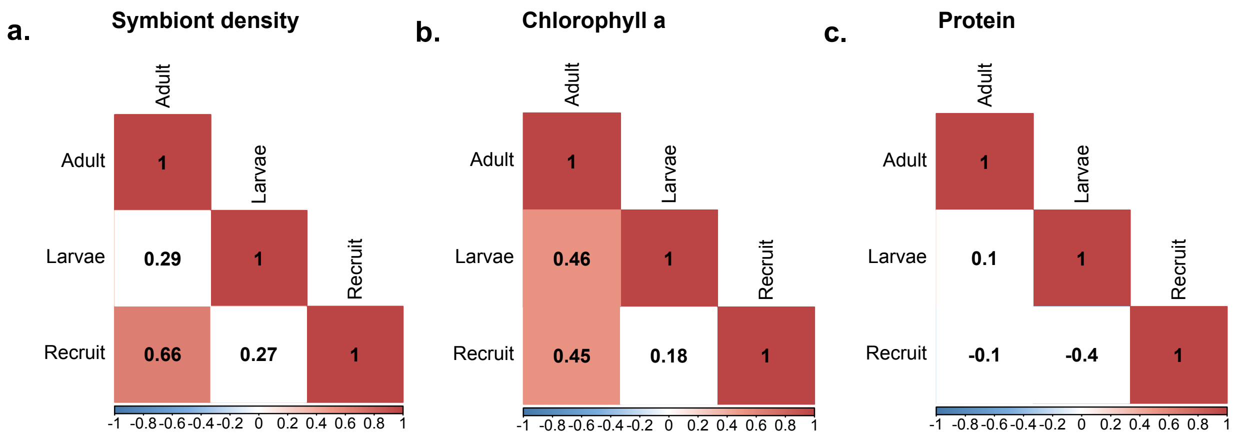


**Figure S1.** Correlation matrices of traits among different life stages. A colored square indicates correlation coefficient for a given comparison is significant ($p < 0.05$).

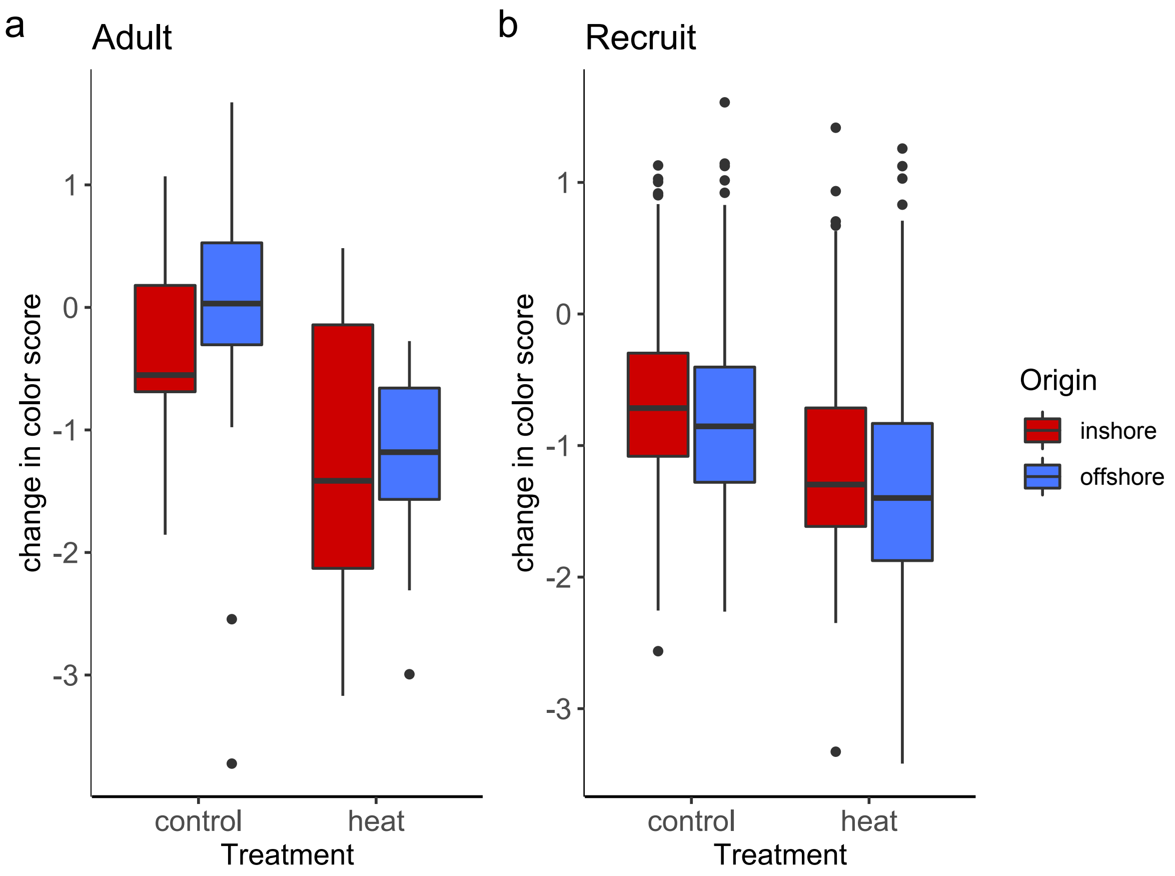


**Figure S2.** Change ($T_{final}$ - $T_{initial}$) in color score for adult and recruit corals (mean ± SEM) in response to experimental conditions separated by reef origin. Positive values indicate increase in pigmentation and negative values indicate decrease in pigmentation (or bleaching).

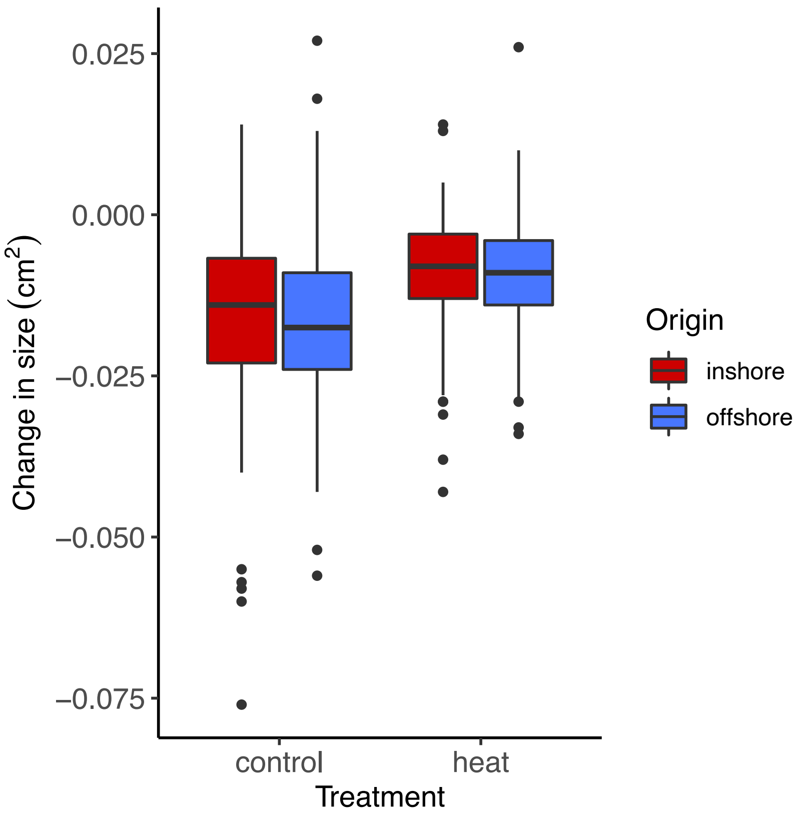


**Figure S3.** Change ($T_{final}$ - $T_{initial}$) in size for recruit corals (mean ± SEM) in response to experimental conditions separated by reef origin.

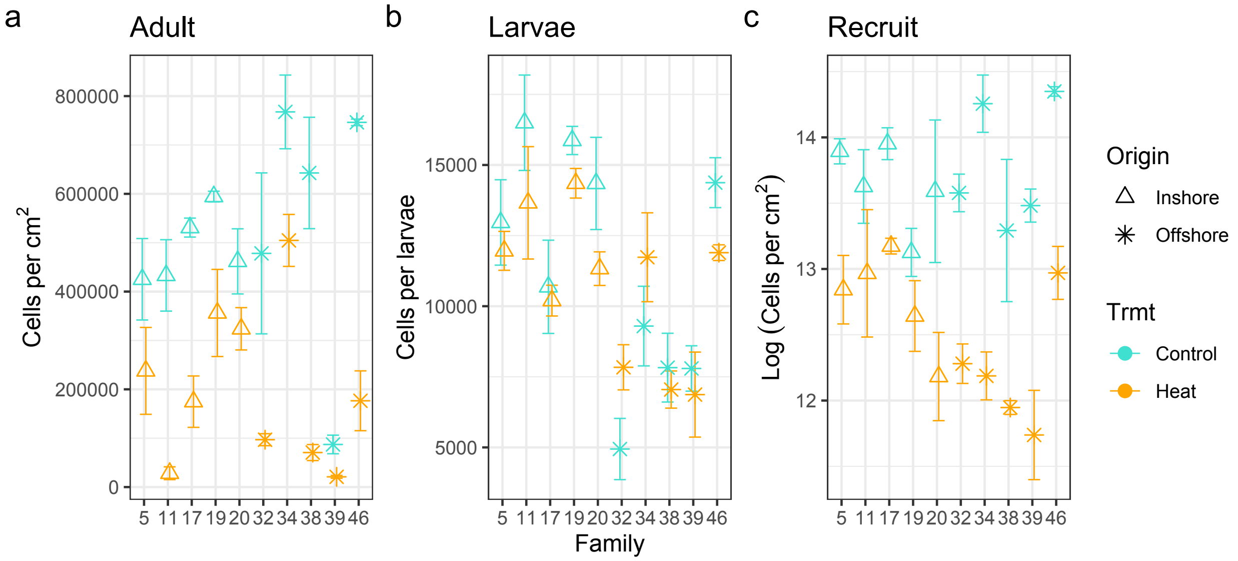


**Figure S4.** Standardized symbiont density (mean ± SEM) of individual *Porites astreoides* adult, larval, and recruit families in response to experimental conditions separated by reef origin. Values were log-transformed if not normally distributed.
